# On the biodegradation of micropatterned polymeric films

**DOI:** 10.1101/2025.01.15.633178

**Authors:** Irene Guerriero, Cristiano Pesce, Raffaele Spanò, Stefania Sganga, Nicola Tirelli, Daniele Di Mascolo, Anna Lisa Palange, Paolo Decuzzi

## Abstract

Polymeric implants for local drug delivery offer significant advantages for treating various medical conditions by enabling the temporal and spatial control of drug release, improving efficacy, and reducing systemic side effects. In this context, µMESH, a 20 μm thin, dual-compartmentalized film comprising a poly(lactic-co-glycolic acid) (PLGA) micronetwork intercalated with a polyvinyl alcohol (PVA) microlayer, represents an interesting opportunity as its geometry can be systematically and accurately micropatterned during the fabrication process, enabling the systematic analysis of the effect of geometry on biodegradation rates mechanisms. In this study, four different µMESH films were realized with different surface area-to-volume ratios (*S*_*a*_/*V*), ranging from 0.67 to 1.7 µm^−1^. After characterizing the µMESH geometry via fluorescent and scanning electron microscopy, biodegradations studies were performed up to 60 days in different media to assess the mass loss of PLGA, the reduction in PLGA molecular weight, and the formation of macroscopic defects – pores, holes and crack – within the PLGA micronetwork. By comparing the four µMESH films among themselves and to a flat, continuous PLGA slab (FLAT), it was confirmed the importance of the surface-to-volume ratio and demonstrated that µMESH with higher *S*_*a*_/*V* ratios exhibited slower degradation rates compared to FLAT. Scanning electron microscopy images of the PLGA micronetworks revealed morphological changes indicative of bulk erosion, including surface roughening and pore formation, in FLAT and µMESH configurations with low *S*_*a*_/*V* ratios. These findings confirm that film micropatterning significantly influences degradation kinetics.

## Introduction

There is growing interest in developing drug delivery systems (DDS) due to their ability to efficiently deliver several therapeutics, including antibodies, peptides, drugs, and enzymes, in a controlled and sustained manner. DDS enable targeted delivery to specific tissues or organs, enhancing drug penetration, reducing the required dosage, and minimizing systemic toxicity [1, 2]. This is achieved through the use of biomaterials, whose properties – such as size, shape, surface chemistry, charge, and morphology – can be tailored to realize DDS with specific release profiles [3]. Among biomaterials, polymers are widely used and investigated. Poly(lactic-*co*-glycolic acid) (PLGA) is a copolymer already used in numerous FDA approved products for human applications [4, 5]. A key advantage of using PLGA is its ability to degrade in aqueous environment through hydrolysis of its ester backbone, generating lactic acid (LA) and glycolic acid (GA) as byproducts. These by-products do not interfere with cells growth and are naturally removed from the body through normal metabolic pathways [5, 6].

PLGA is classified as a bulk eroding polymer, with water uptake is faster than the rate of hydrolysis [6, 9]. Hence, the degradation process occurs uniformly across the polymeric matrix until a critical molecular weight is reached. The prolonged retention of acidic by-products leads to localized acidification within the matrix, which in turn promotes the ester hydrolysis through an autocatalytic effect [10]. The degradation rate of PLGA is influenced by several factors, including the molecular weight, length and ratio of lactic and glycolic blocks, as well as pH value, temperature, and buffering capacity of the surrounding medium [6]. Furthermore, the shape and size of the polymeric device influence the degradation process, with the surface area-to-volume ratio (SVR) playing a critical role in diffusion-limited processes such as ester hydrolysis [3]. Indeed, several studies investigated the ability to modulate the degradation velocity and mitigate the bulk erosion mechanisms by changing the dimensions of micro- and nano-particles, with faster degradation rates being associated with larger dimensions [11].

In this context, µMESH serves as an interesting example of a thin conformable, dual-compartment film consisting of a PLGA micro-network integrated with a micropatterned polyvinyl alcohol (PVA) microlayer. The µMESH architecture enables the encapsulation of hydrophilic and hydrophobic drugs inside the PVA and PLGA compartments, respectively [7, 8]. While PVA dissolves quickly in aqueous environments releasing its therapeutic or imaging cargo, drug release from the PLGA micro-network is regulated by both drug diffusion and polymer degradation [3, 9]. Since the fabrication process is versatile and allows for the fine-tuning of the geometric features, resulting in µMESH with different micro-metric dimensions, this DDS provides the opportunity to characterize rates and mechanisms of biodegradation for micropatterned PLGA films of different geometries. In this work, four different micropatterned µMESH films have been realized and compared to a flat, continuous layer of PLGA (FLAT) to investigate the effect of the architecture and surface-to-volume ratio on biodegradation rates.

## Materials and Methods

### Materials

Dimethylsulfone (DMSO_2_, TraceCERT, 99.99%), dimethyl sulfoxide anhydrous (DMSO), dimethyl sulfoxide-d6 99.9 atom % D ((CD3)2SO), Float-A-Lyzer (cut-off 100 kDa), poly(D,L-lactide-co-glycolide) (lactide/glycolide 50:50, Mn 38,000−54,000), poly(vinylalcohol) (PVA, Mw 9000−10,000 Da, 80% hydrolyzed), Gill No.2 Haematoxylin solution (Sigma-Aldrich, GHS232), Eosin Y alcoholic solution (Sigma-Aldrich, HT110132), sucrose (Sigma-Aldrich, S7903), and sodium azide were purchased from Merck (Darmstadt, Germany). Curcumin (95% total curcuminoid content) from Turmeric rhizome and trifluoroacetic acid (TFA) 99% were purchased from Alfa Aesar GmbH & Co KG (Karlsruhe, Germany). Tubes used for the NMR analysis were disposable for SampleJet NMR with 3 mm O.D. and were purchased from Bruker Italia Srl (Milan, Italy). Paraformaldehyde solution 4% in PBS (product code sc-281692) was from Santa Cruz Biotechnology Inc. (Heidelberg, DE). Surgipath® FSC 22 Frozen Section Compound was purchased from Leica Microsystems GmbH (Wetzlar, DE).

### µMESH fabrication

µMESH is prepared following a soft lithographic method, as previously described by the authors [8]. Briefly, direct laser writing was used to engrave the main geometric features of µMESH on a silicon template. Specifically, 5 µm deep and 3 µm wide channels describe a regular array of squared pillars, with edge lengths of 5, 10, 20, and 50 µm. It is noteworthy that all the geometric parameters can be easily modified by reprogramming the direct laser writing steps. This micropatterned geometry of the silicon master template was replicated using a molding process. First, a PDMS layer was obtained by pouring the polymeric base and the curing agent (10:1 ratio) on top of the micropatterned silicon master and drying this mixture at 60°C for 4 hours. Then, the PDMS template was used to further replicate the geometry into a PVA microlayer, obtained by pouring the polymeric solution over the PDMS template and drying it at 60°C for 2 hours. Upon complete water evaporation, the resulting PVA microlayer was peeled off from the PDMS template. The PVA microlayer retains the identical geometrical pattern as the original silicon template. The resulting channels (ridges) were filled with a polymeric paste made by dissolving known amounts of PLGA in acetonitrile (ACN). The resulting 30 mm square composite film comprises the PVA microlayer (hydrophilic compartment) and the PLGA micronetwork (hydrophobic compartment). This composite film was then cut into multiple 5 mm square pieces, referred to as µMESH. Additionally, a composite film without any geometric pattern was fabricated by spreading the PLGA polymeric paste over a flat PVA layer, resulting in a bi-layer composite film referred to as FLAT.

Hydrophobic molecules can be easily incorporated into the PLGA paste. Specifically, 500 µg of green, fluorescent curcumin (CURC) were added per template to assess loading efficiency. Confocal microscopy (A1^+^/A1R^+^, Nikon corporation, Tokyo, Japan) was also employed to image the resulting µMESH and FLAT samples, confirming the successful formation of the PLGA hydrophobic micro-network.

### µMESH biodegradation

To evaluate µMESH biodegradation over time, the PLGA micronetwork was first separated from the PVA microlayer. Note that given the molecular weight (9,000 – 10,000 Da) and rate of hydrolysis (80%) considered in this work, the PVA microlayer dissolves rapidly following exposure to a physiological solution. Therefore, the dual-compartmentalized µMESH was placed into a Petri dish with the PVA component facing downwards. Then, DI water was dropped on the airside. After 2 hours at RT, the remaining PLGA micronetwork was gently washed to ensure full detachment from the substrate. The samples were swabbed with blotting paper, transferred on a nylon sheet, and dried at RT.

The collected PLGA micronetworks were placed inside a sealed jar filled with 50 ml of DI water or 0.1 M PBS pH 7.4 containing 0.02% sodium azide. The number of micronetworks was adjusted to ensure that all the jars contained the same initial amount of PLGA (15 ± 0.05 g). The jars were placed into an incubator at 37.0 ± 0.1°C, and samples were taken at pre-determined time points to monitor degradation, namely 7, 14, 21, 28, and 60 days. To avoid affecting the erosion mechanisms, the degradation media were not replaced during the study.

### Mass loss measurements

Samples were weighed before being placed in the degradation medium to determine the initial mass of the PLGA micronetwork, *m*_*ini*_. After fixed time intervals, the PLGA micronetoworks were recovered by filtration, using a nylon sheet with a pore size of 40 µm (pluriSelect life science UG & Co. KG, Germany). The samples were gently washed with DI water and dried at RT to record the dry mass (m_dry_) left after degradation. The mass loss of the PLGA micronetworks was calculated as:

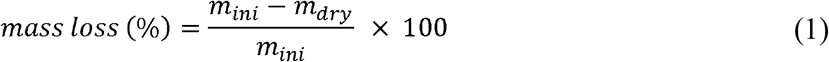

The PLGA micronetworks were recovered at predetermined time intervals to monitor degradation until the mass loss reached approximately 75% of the initial mass *m*_*ini*_, on average.

### Potentiometry

pH measurements were performed with a glass-bodied combination pH electrode (InoLab pH730, WTW GmbH, Weilheim, Germany). The pH-meter was calibrated with three buffer solutions: buffer solution pH 4.01, phosphate buffer solution pH 7.00, and buffer solution pH 10.01. After recovering the PLGA micronetworks from the jars and the dialysis devices from the incubation tubes, the pH value of the degradation media was recorded.

### Gel permeation chromatography

The number average and weight average molecular weights 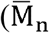 and 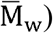 were determined by gel permeation chromatography (GPC). Samples were recovered at predetermined time points, namely 7, 14, and 21 days, and processed. First, samples were freeze-dried, then dissolved in N,N-Dimethylformamide (DMF) with 2.5 mg/l LiBr at 50°C, at a concentration around 10 mg/mL for 8 hours, and finally filtered through a 0.22 μm PTFE filter. GPC was performed using an integrated OMNISEC system (Malvern Panalytical Ltd., UK) equipped with a D6000M and a D2500 column (10 and 6 μm particle size, both 300 × 8 mm) and a triple detection method (refractive index, a viscometer, and a dual angle light scattering detector at 15° and 90°). DMF with 2.5 mg/l LiBr was used as an eluent at a temperature of 50 °C and a flow rate of 1 ml/min. The system was calibrated using poly(methyl methacrylate) (PolyCal standards, Malvern Panalytical Ltd., UK) 51 kDa narrow standard with known dispersity, intrinsic viscosity, and dn/dc. For each condition, measurements were averaged, and the 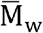 values were plotted against time. Points were fitted using a linear equation, with slope calculated as:

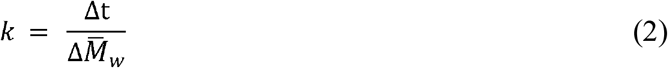

where *k* describes the steepness and direction of fit.

### PLGA quantitative analysis by q-NMR

First the PVA microlayer was removed from µMESH (*n* = 10) by carefully rinsing, drying and transferring the remaining PLGA micronetwork samples into vials containing 2 ml of DI water or 0.1 M PBS pH 7.4. The vials were incubated at 37 ± 0.1°C. At predetermined intervals, namely 0, 7, 14, 30, and 60 days, samples were collected and centrifuged to separate any soluble degradation products (12,700 rpm, 4°C, 30 min). After replenishing the supernatant with fresh DI water, a second centrifugation was performed. The resulting pellets were freeze-dried, and the lyophilized samples were dissolved in a known volume of 3% trifluoroacetic acid (TFA) in (CD_3_)_2_SO and then loaded into 3 mm disposable SampleJet tubes. All NMR spectra were recorded on a Bruker Avance III 400 MHz spectrometer. Quantitative analyses were carried out automatically with 16 scans, 65536 data points, and an inter-pulse delay of 40 s over a spectrum of 20.49 ppm (offset at 6.17 ppm). Spectra were manually phased and automatically baseline corrected.

The so determined PLGA concentration was normalized based on the degree of polymerization (*DP*_*ave*_), approximately 353.6. This value was calculated following the equation:

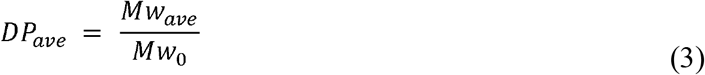

*w*here *Mw*_*ave*_ corresponds to the average molecular weight of the polymer chain, ranging between 38 and 54 kDa, and *Mw*_0_ to the molecular weight of the individual repeating units, which is 130.1 Da.

### Scanning electron microscopy characterization

For each micropattern, one 5 mm square µMESH was placed into a 35×10 mm Petri dish filled with 3 ml of DI water or 0.1 M PBS pH 7.4 and incubated at 37.0 ± 0.1°C. Before contact with an aqueous solution and at predetermined time points, namely 7, 14, 30 and 60 days, µMESH was taken out from the medium, carefully washed with 3 ml of DI water, and dried at RT. To monitor topographical and morphological changes over time, samples were sputter-coated with 10 nm of gold and then imaged at 10 keV using an analytical low-vacuum SEM observation (SEM □ JSM-6490, JEOL Ltd., Tokyo, Japan).

### Statistical Analyses

To compare two groups, an F-test was performed to verify the absence of statistical significantly different variances, followed by a two-tailed t-test. For multiple comparison (i.e., experiments involving three or more groups), the Brown-Forsythe test was used to assess variance equality across groups, followed by a one-way analysis of variance (ANOVA). The two-sided log-rank test was applied to determine statistical significance. All statistical analyses and graphs were generated using GraphPad PRISM. Results were considered statistically significant for p-values lower than 0.05.

## Results

### µMESH fabrication and characterization

µMESH is a dual-compartmentalized polymeric system consisting of a PVA microlayer intercalated with a PLGA micronetwork (**Figure 1A-B**). It can be loaded with various therapeutic agents, including small molecules, chemotherapeutic drugs, and nanomedicines [7, 8]. µMESH enables the local, sustained release of these agents over periods ranging from several days to months, making it particularly suited for the treatment of high-grade gliomas using potent drugs that, if administered systemically, would not accumulate in the brain at sufficient levels (poor brain penetrance) and cause intolerable side effects. The archetypal configuration of µMESH – the 20×20 µMESH with square openings of 20 µm – has demonstrated therapeutic efficacy in preclinical animal models of orthotopic glioblastoma [7, 8] Additionally, given the modularity of the fabrication process, µMESH can be realized with different geometries tailoring its physio-chemical and pharmacological properties based on the intended application.

**Figure 1:**
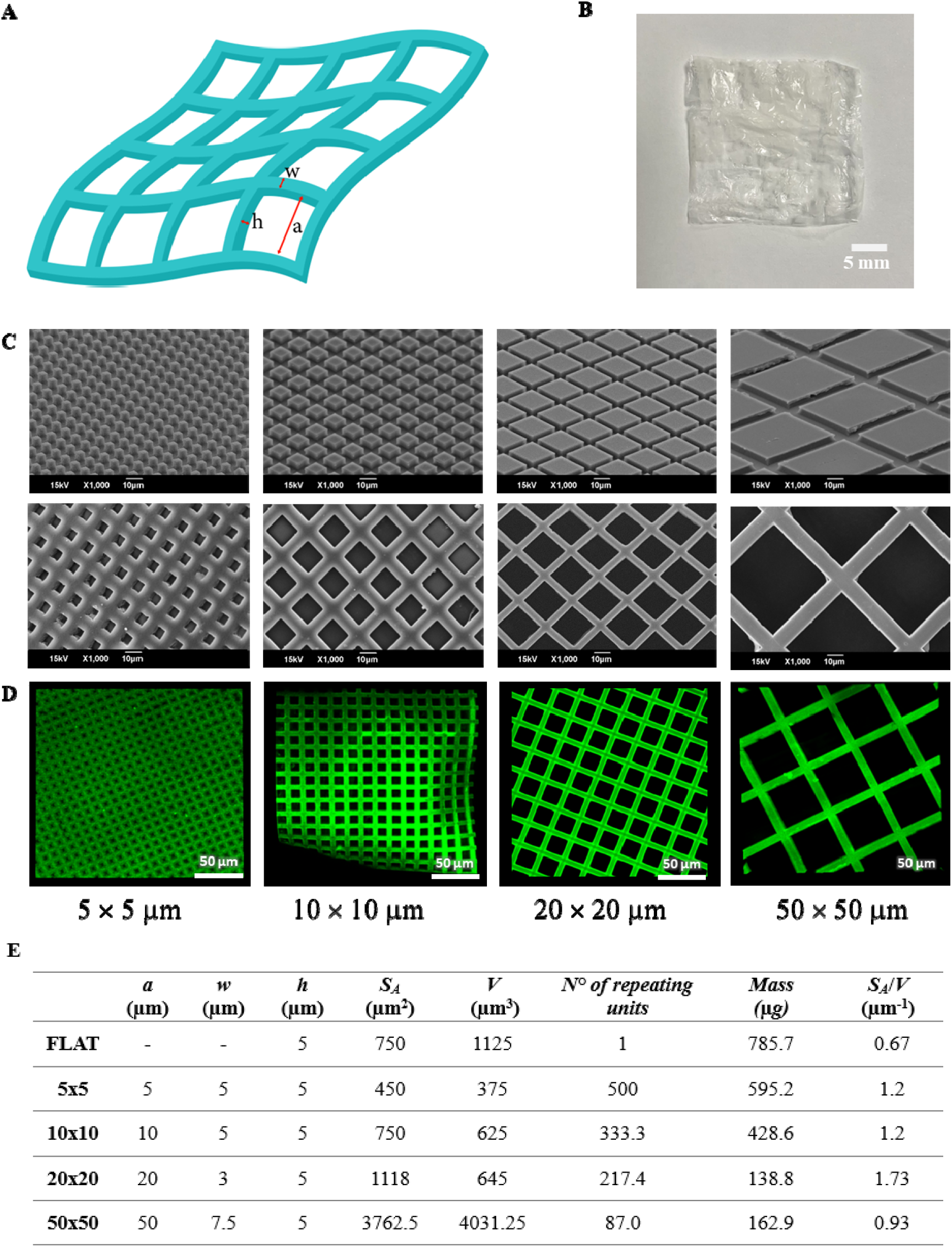
Different µMESH configurations. **A.** Schematic representation of µMESH highlighting its geometrical attributes. **B**. Optical image of µMESH after PVA dissolution in aqueous solution. **C**. Scanning electron microscopy images of the PVA microlayer (***first row***) and the PLGA micronetwork (***second row***), after PVA dissolution in aqueous solution. **D**. Confocal fluorescent microscopy images of the PLGA micronetwork, after PVA dissolution in aqueous solution. **E**. Table listing the geometrical properties of four µMESH configurations (5×5, 10×10, 20×20, and 50×50 µMESH) and FLAT.

As detailed in the **Methods** section, µMESH is fabricated through a top-down approach. The first lithographic step is crucial for accurately defining the final geometry of µMESH, as it determines the size and shape of an array of micropillars on a master silicon template by adjusting the direct laser writing parameters. The designed micropattern is then transferred into a PVA microlayer via an intermediate PDMS template. The PVA microlayer replicates the exact geometrical features of the silicon master. In this work, four specific µMESH configurations were considered, resulting in PVA microlayers displaying regular arrays of square micropillars with an edge length of 5, 10, 20, and 50 µm (**Figure 1C** – first row). Then, a PLGA solution was carefully applied over the PVA microlayers to uniformly and completely fill the empty ridges within the square micropillars. Upon solvent evaporation, this led to the formation of PLGA microlayers (**Figure 1C** – second row). Confocal fluorescent images of the PLGA microlayers, loaded with the green fluorescent molecule curcumin, are shown in **Figure 1D**. The distance between adjacent PVA micropillars (i.e., the width *w* of the PLGA strands in **Figure 1A**) and the height of the PVA micropillars (i.e., the thickens *h* of the PLGA strands in **Figure 1A**) were adjusted according to the edge length of the PVA micropillars (i.e., the size *a* of the square openings in the PLGA micronetwork, as in **Figure 1A**). The table in **Figure 1E** summarizes the geometrical properties of the µMESH configurations. Using the archetypal 20×20 µMESH as a reference and having fixed the thickness *h* of the PLGA strands at 5 µm, the width *w* of the PLGA strands increases to 7.5 µm for the 50×50 µMESH to ensure adequate mechanical strength and to 5 µm for the 10×10 and 5×5 µMESH to allow the reduction in size of the square openings. This results in µMESH configurations with varying surface areas *S*_*a*_ and volumes *V* of the PLGA micronetwork. Specifically, the surface area *S*_*a*_ of one, single unit in 20×20 µMESH was ∽1,100 µm^2^, decreasing to 750 µm^2^ for the 10×10 µMESH and 450 µm^2^ for the 5×5 µMESH, while it increased nearly fourfold for the 50×50 µMESH. The volume *V* filled with PLGA was ∽650 µm^3^ for the 20×20 and 10×10 µMESH, almost doubling in volume for the 5×5 µMESH, and nearly sixfold for the 50×50 µMESH. The values associated with the 5 mm square µMESH are obtained considering the total number of repeating units (openings) (**Figure 1E**). Consequently, the surface area-to-volume ratio (*S*_*a*_/V) changed too with the µMESH configurations, being ∽1.7 for the 20×20 µMESH, reduced to 1.2 for both the 10×10 and 5×5 µMESH, and to 0.93 for the 50×50 µMESH.

In the following, the various µMESH configurations are compared to FLAT, which consists of an uncarved, flat PVA microlayer onto which a PLGA layer was uniformly deposited. FLAT represents a conventional polymeric implant with no openings.

### µMESH biodegradation assays

The geometry of µMESH is expected to influence its biodegradation process, as well as its pharmaceutical and biological performance. To determine the correlation extent between geometrical parameter – *S*_*a*_/V – and degradation behavior, a first investigation was conducted by fixing the total mass of PLGA mass.

Previous studies have shown that PVA can influence the degradation rates of PLGA micro- and nanoparticles [10, 11], thus the degradation rates under different conditions were assessed by first removing the PVA microlayer and then monitoring the long-term dissolution of the PLGA micronetwork. It is here important to recall that the dissolution of PVA in water depends on factors such as the degree of hydrolysis and molecular weight. A high degree of hydrolysis and high molecular weight favor interactions between PVA molecules over those between PVA and water, thereby reducing the dissolution rate [12]. In this work, the supporting microlayer was fabricated using PVA with a relatively low degree of hydrolysis (80%) and a low molecular weight (9 – 10 kDa), ensuring dissolution within a few minutes upon exposure to water.

After placing platforms in DI water for 2 hours, the remaining PLGA micronetwork was swabbed with paper and dried at room temperature, resulting in opaque film as shown in **Figure 1B**. Then, the hydrolytic degradation was assessed by measuring the loss in weight over the course of 60 days upon incubation in DI water and PBS (pH 7.4) (**Figure 2A** and **2B**, respectively), at 37°C. Specimens were incubated without stirring, since body fluids move slowly in solid tissues, which are the typical implantation site of µMESH. Similarly, the pH variation of the medium was monitored over time (**Figure 2C**). Note that the number of µMESH per experiment was adjusted to have the same initial mass of PLGA (15 ± 0.05 g) in solution for all tested configurations.

**Figure 2:**
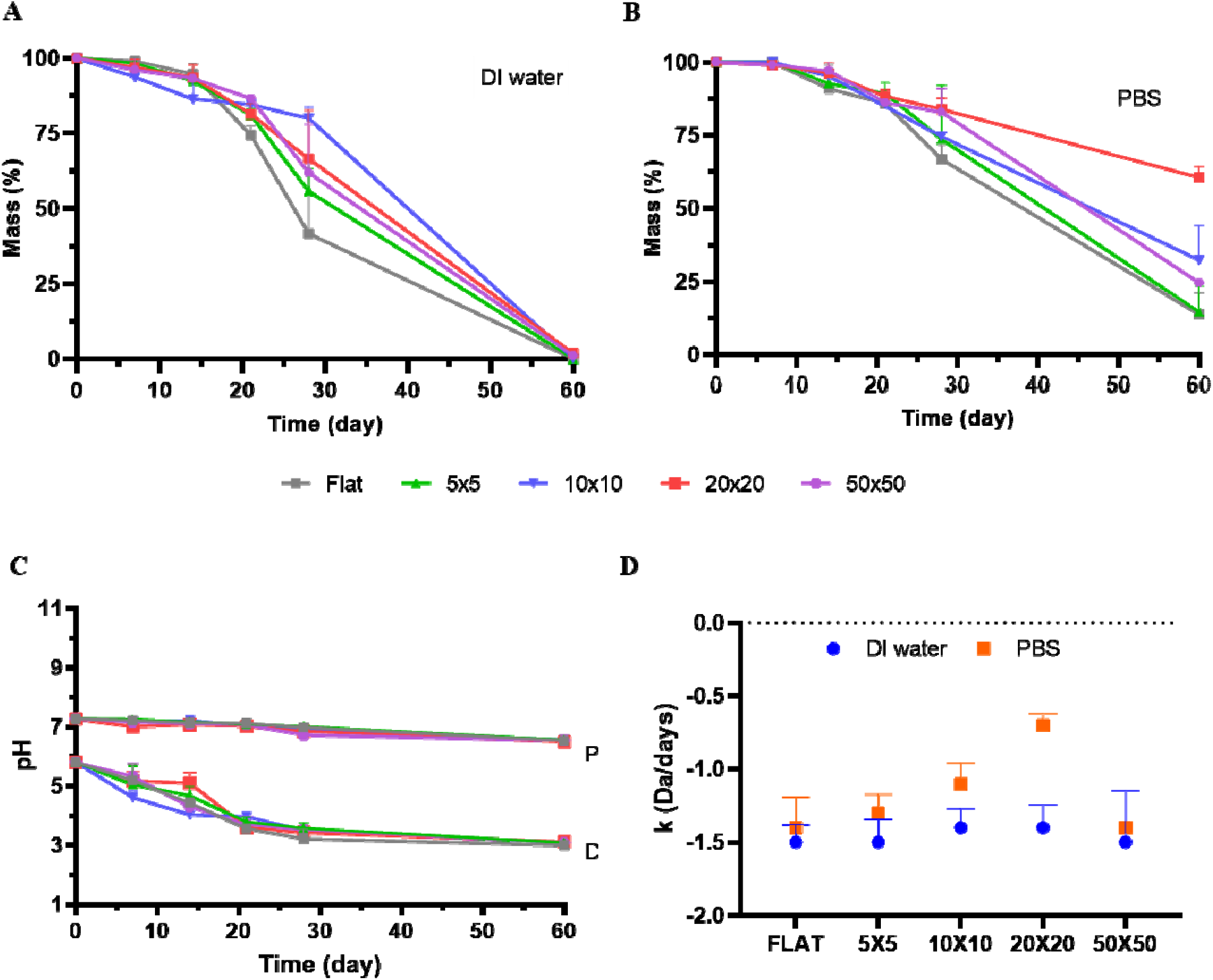
µMESH biodegradation. **A. – B**. mass loss versus time for the PLGA micronetworks in DI water and PBS for all tested µMESH configurations and FLAT. **C**. Change in pH versus time for the PLGA micronetworks in DI water and PBS for all tested µMESH configurations and FLAT. **D**. Rate of reduction for the molecular weight of the PLGA micronetwork in DI water and PBS, for all tested µMESH configurations and FLAT, via gel permeation chromatography.

In DI water, mass loss was found to be lower than 10% for all µMESH configurations and FLAT during the first 7 days of observation (**Figure 2A**). From day 7 onward, mass loss occurred more rapidly with a significant variation observed on day 21 among the different tested µMESH configurations and FLAT. The 10×10 and 20×20 µMESH had the lowest mass loss, corresponding to approximatively 20% and 30%, respectively. They were followed by the 5×5 and 50×50 µMESH that displayed a mass loss of nearly 40%. The FLAT showed the fastest degradation with a mass loss of 60% on day 21. At the endpoint, 60 days, all the tested µMESH configurations and FLAT were fully degraded with no residual mass left. Interestingly, a physical examination of the samples revealed that the PLGA micronetwork softened immediately upon immersion in DI water, suggesting rapid water diffusion into the PLGA strands. As the immersion continued, the micronetwork became progressively cloudy and whitish, indicative of a change in light refractive index. Once dried, all the specimens shrunk in their dimension **(Supporting Figure 1**) and their brittleness increased with the incubation time. At later time points, dried residual PLGA micronetworks appeared as a compact, viscous mass.

In PBS, the mass loss is again nearly zero over the first 7 days, but it increases quasi linearly over the following 7 weeks of observation (**Figure 2B)**. Among all the µMESH configurations, the 20×20 µMESH had the lowest mass loss, corresponding to approximatively 10% on 21 days and just 40% at 60 days. This was followed by the 10×10 and 50×50 µMESH with mass losses at 60 days of approximatively 70%, while 80% of the mass was degraded for the 5×5 µMESH. Even in a physiological solution, FLAT presented the fastest degradation with a mass loss of 80% on day 60. In strike difference with the DI water results, residual portions of µMESH were observed at the end of the experiment. PBS is commonly used as an acceptance medium in biodegradation and drug release studies due to its ability to maintain pH and osmolarity consistent with body fluids [6]. However, to more accurately mimic *in vivo* conditions, the degradation of a 5×5 μMESH was also performed in artificial cerebrospinal fluid, showing a comparable trend in mass loss and pH variation to that observed in PBS (**Supporting Figure 2**).

**Figure 2C** gives the change in pH during the degradation of the PLGA micronetwork. It is well known that PLGA hydrolysis is associated with the local acidification of the solution. Indeed, this is evident for the studies conducted in DI water where the pH progressively decreased from about 5.9 ± 0.10 on day 0 to 3.0 ± 0.10 on day 60, for all tested µMESH configurations and FLAT, with small variations within the first 3 weeks. However, no significant change in pH was documented for the experiments conducted in a physiological solution where the pH progressively decreased from about 7.2 ± 0.10 on day 0 to 6.5 ± 0.10 on day 60, for all tested samples. Comparable results were obtained in a different assay where µMESH and FLAT, including the PVA microlayer, were allowed to degrade within Float-A-Lyzer dialysis devices (**Supporting Figure 3**).

The rate of degradation of the PLGA micronetwork was further verified by monitoring changes in its molecular weight over time using GPC. Following incubation in DI water and PBS, the rate of reduction *k* of the PLGA molecular weight is depicted in **Figure 2D**, for all tested µMESH configurations and FLAT. In agreement with the mass loss analyses, the samples incubated in DI water experienced a much more rapid degradation, as documented by the lower *k* values compared to PBS. Once again, the 20×20 and 10×10 µMESH configurations presented the lowest degradation rates, as demonstrated by the larger *k* values, both in DI water and PBS. A negligible difference was observed in the eroding kinetics for FLAT and 50×50 µMESH when incubated in DI water and PBS.

Finally, µMESH biodegradation was assessed *via* NMR. Note that in this experiment, the number of µMESH, rather than the mass of PLGA, was fixed. µMESH presents a macroscopic superficial square area of 5 mm, thus the varying micropattern is reflected in both *S*_*a*_/V value and amount of PLGA (as reported in the Table in **Figure 1E**). Specifically, the 50×50 geometry is associated with a lower *S*_*a*_/V and a lower PLGA mass than the ones associated with the 5×5 and 10×10.

Ten µMESH for each configuration and FLAT were incubated in DI water and PBS within a Float-A-Lizer dialysis device. At predetermined time points, samples were withdrawn from the medium, and the residual contents of PVA and PLGA were analyzed using a protocol based on PULCON method (**Supporting Information, Supporting Figure 4** and **5**), previously described by the authors [10]. qNMR spectra were then generated for the different µMESH configurations and FLAT, at predetermined time points and after incubation in different media. PVA microlayer rapidly dissolved, losing ∼98% of its original mass within the first hour of incubation. As previously reported, the PVA microlayer rapidly dissolved, losing ∼98% of its original mass within the first hour of incubation [13]. To quantify the mass of PLGA in the micronetwork, the peak at 5.05 – 5.30 ppm, corresponding to the methine (-CH-) functional group (**Supporting Figure 5**), was considered. In **Figure 3A**, showing one representative qNMR spectra, this peak appears well isolated from the others, thus minimizing measurements inaccuracies. **Figure 3B** and **3C** illustrate the variation in polymer mass over time, as measured via the qNMR PULCON protocol. Since the number of µMESH was fixed, the parametric curves start from different initial masses. In both DI water (**Figure 3B**) and PBS (**Figure 3C**), the FLAT exhibited the largest change in mass over the 60-day observation period. In PBS, the average mass loss rate, calculated as the ratio between the total mass loss and the duration of the observation period, was 3 µg/day for the FLAT, followed by 2.25 µg/day for the 5×5 µMESH, 1.67 µg/day for the 10×10 µMESH, 0.83 µg/day for the 50×50 µMESH, and 0.67 µg/day for the 20×20 µMESH. In this assay, a significant difference in the degradation rate of the 50×50 µMESH was observed compared to the degradation data collected in **Figure 2**, where the 5×5 and 10×10 µMESH exhibited a similar behavior.

**Figure 3:**
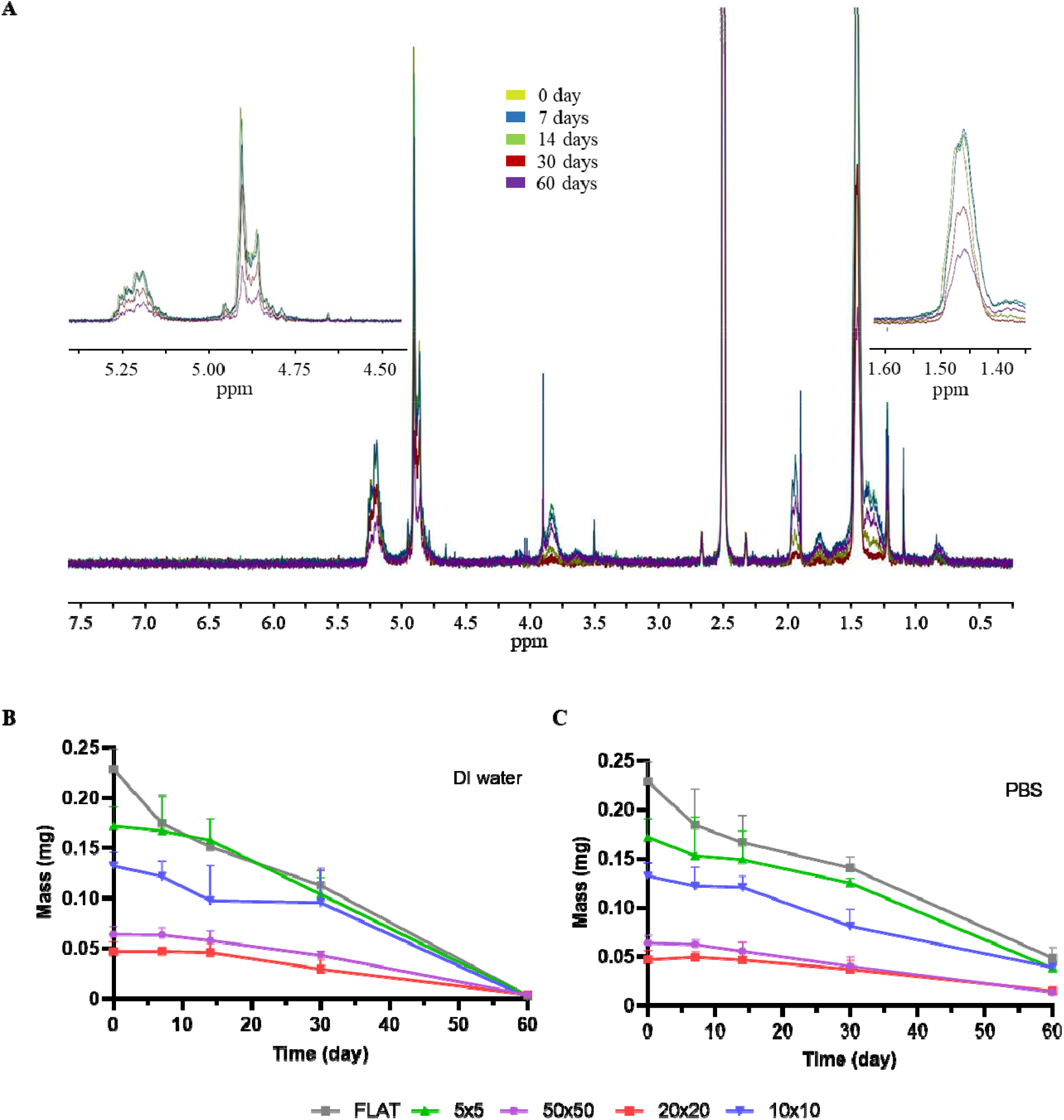
µMESH biodegradation via qNMR analysis. **A**. 1H *q*NMR spectra of a 20×20 µMESH incubated in PBS at predetermined time points, over a 60-day observation period. **B**. Mass loss vs. time for µMESH in DI water for all tested µMESH configurations and FLAT, via the qNMR PULCON protocol. **C**. Mass loss vs. time for µMESH in PBS for all tested µMESH configurations and FLAT, via the qNMR PULCON protocol.

### µMESH morphological alternations over time

Changes in the µMESH morphology over time were evaluated in DI water and PBS using scanning electron microscopy (SEM), as summarized in **Figure 4**. Starting from day 0, each row presents SEM images for the four different µMESH configurations and FLAT, alternating in DI water and PBS at predetermined time points. Surface roughening and the formation of pores, holes, and cracks were considered as key indicators of the degradation process. In DI water, µMESH started to exhibit these degradation marks as early as 14 days, with their size and abundance increasing over time. By 30 days of incubation, all µMESH configurations had lost their original geometry, with PLGA strands bundling together to form a porous solid structure. By 60 days, the structures had shrunk and collapsed, closing the pores and holes to form a continuous, solid, irregular layer. In contrast, µMESH incubated in PBS retained their distinct shape up to 60 days. However, the PLGA strands appeared thinner in the 20×20 and 50×50 µMESH configurations.

**Figure 4.**
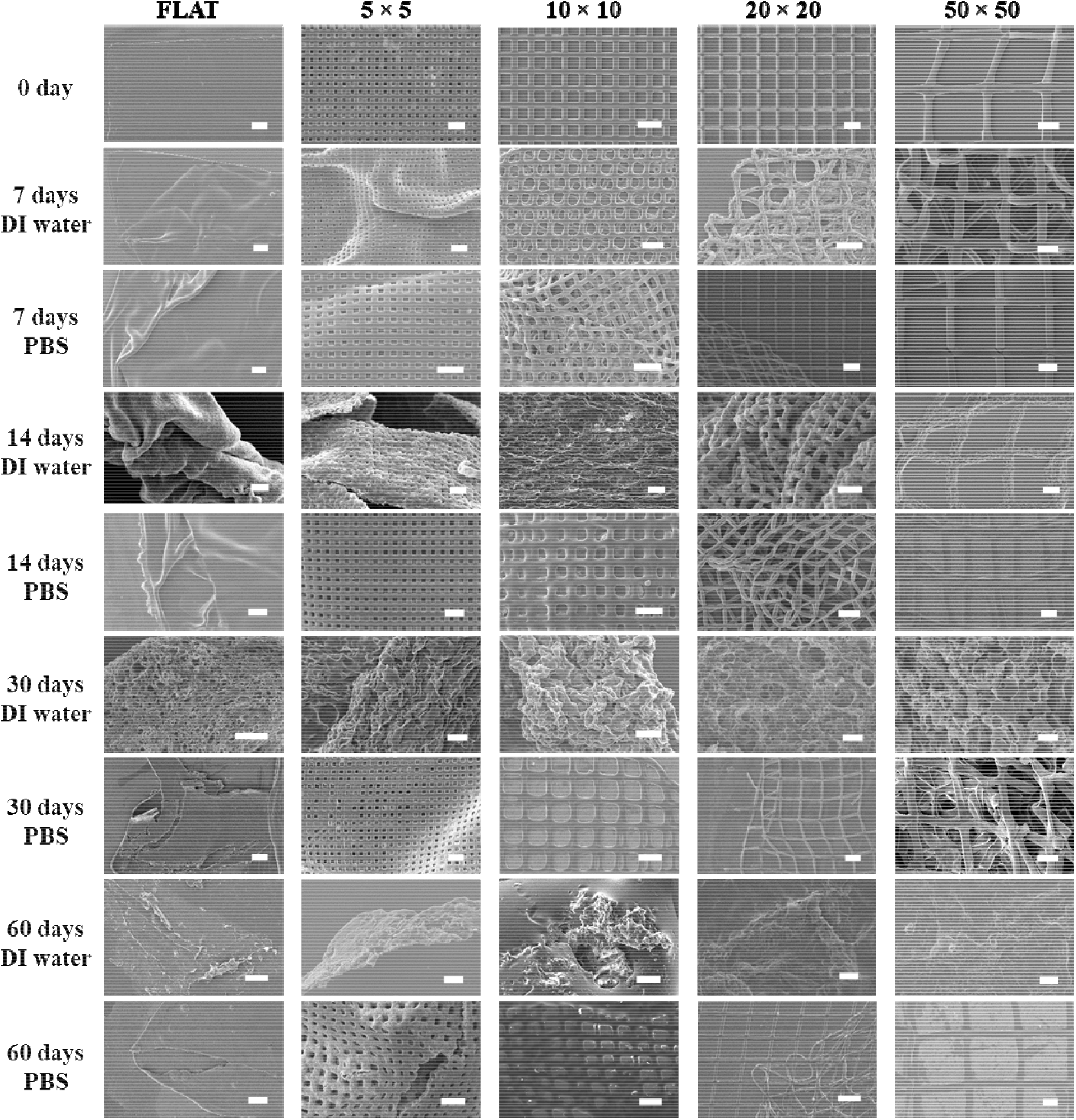
µMESH morphological alternations over time. Morphological alternations for all tested µMESH configurations and FLAT after 7, 14, 30, and 60 days in DI water and PBS. From left to right: FLAT (control); 5×5 µMESH; 10×10 µMESH; 20×20 µMESH; and 50×50 µMESH (Scale bar: 100 µm for Flat system, and 20 µm for 5×5, 10×10, 20×20 and 50×50 geometry).

## Discussion

In prior studies, the 20×20 μMESH was used to treat orthotopic murine models of glioblastoma with chemotherapeutic and anti-inflammatory drugs loaded into the PLGA hydrophobic micronetwork [7, 8]. In this study, four µMESH configurations were fabricated using a versatile replica-molding technique leading to PLGA micronetwork with varying sizes of the square openings and widths of the polymer strands, as summarized in the table of **Figure 1E**. These enable the generation of PLGA micronetworks with different *S*_*a*_ */V* ratios: FLAT and the 50×50 µMESH had the lowest *S*_*a*_ µMESH had the highest *S*_*a*_*/V*, being equal to 0.67 and 0.93 µm^−1^, respectively; the 20×20*/V* µMESH had the highest *S*_*a*_*/V* = 1.73 µm^−1^; the 5×5 and 10×10 µMESH shared intermediate values of *S*_*a*_*/V* = 1.2 µm^−1^. Therefore, the impact of the µMESH micropatterned geometry on biodegradation was compared both among four different µMESH configurations and with an unpatterned composite microlayer (FLAT), represented by a 5 µm thick solid PLGA slab.

PLGA degrades *via* hydrolysis, breaking ester bonds into lactic acid and glycolic acid [11]. Hydrolysis occurs when water penetrates the polymer matrix, with two primary mechanisms: bulk erosion and surface erosion. In bulk erosion, water permeates the matrix faster than it degrades, causing a uniform hydrolysis throughout the core. This leads the acidic byproducts to progressively accumulate in the core, lowering the local pH and accelerating degradation through autocatalysis. Bulk erosion typically exhibits an initial lag phase with minimal weight loss, followed by rapid degradation, resulting in matrix softening, structural collapse, and loss of mechanical stability [12] [6, 13]. In surface erosion, water permeation in the matrix is slower than it degrades, confining hydrolysis to the surface. This maintains the integrity of the polymer matrix for longer and the erosion rate is constant and determined by the interaction of water molecules with the surface. Various studies have documented that degradation rates in polymeric scaffolds, microparticles, and nanoparticles strongly depend on the surface-to-volume ratio rather [3, 14] [6]: low *S*_*a*_ */V* ratios slow water diffusion and trap acidic byproducts in the core, enhancing autocatalysis and accelerating degradation; whereas high *S*_*a*_ */V* ratios promote the diffusion of acidic byproducts, reducing autocatalysis and favoring surface erosion. Therefore, it was hypothesized that µMESH degradation and release kinetics would increase with higher *S*_*a*_ */V* ratio.

*As* µMESH with different microgeometry differ both in *S*_*a*_ */V* ratio and PLGA mass and the two might exert a comparable influence on biodegradation behavior, a first investigation was conducted adjusting the number of polymeric micronetworks to have the same initial total mass of PLGA and evaluating mass loss and molecular weight reduction over time.

In DI water, where buffering is absent, all µMESH configurations and FLAT showed faster degradation and molecular weight reduction rates, thereby inducing a rapid pH decrease, which reached values as low as 3 at day 60. In contrast, in PBS and artificial cerebrospinal fluid, the drop in pH was modest from 7.4 to 6.5 after 60 days of incubation, confirming the effect of the media on degradation processes. The 5×5 µMESH (*S*_*a*_*/V* =1.2 µm^−1^) and the 50×50 µMESH (*S*_*a*_*/V* =0.67 µm^−1^) showed higher molecular weight reductions than the 10×10 µMESH (*S*_*a*_ */V* =1.2 µm^−1^), which exhibited intermediate molecular weight reduction rates. Despite identical surface-to-volume ratio, the 10×10 µMESH had a much larger surface area than the 5×5 µMESH, which likely favored escape of acidic byproducts (lactic and glycolic acid) and reduced autocatalysis. However, in PBS, the average reduction in molecular weight appeared to be slower for geometry associated with higher *S*_*a*_ */V* values.

*A*ll these findings suggest a clear dependence of the degradation rate on geometrical features in a linear fashion.

Then, the biodegradation of µMESH was assessed under the initial condition of fixing the total number of polymeric micronetworks via qNMR. The amount of residual PLGA mass over time decreased while moving from the FLAT to the 5×5 and 10×10 µMESH, and eventually to the 50×50 and 20×20 µMESH, unrevealing the influence of the different starting PLGA amount. Specifically, FLAT has the highest mass of PLGA (785.7 µg) while the 20×20 µMESH has the lowest one (138.8 µg). Therefore, both *S*_*a*_ */V* and polymer mass affect the degradation properties of µMESH. The 20×20 µMESH, with its high *S*_*a*_ */V* ratio, exhibited a slower degradation rate, as further confirmed by SEM imaging. In particular, the SEM analysis showed a gradual thinning of the polymer strands and expansion of square openings in the PLGA micronetwork without significant architectural collapse over 60 days in PBS. In contrast, FLAT, with its low *S*_*a*_ */V* ratio, degraded fastest, developing cracks and pores, followed by structure collapse into an aggregated, unstructured mass, which are clear signs of bulk erosion [6, 13]. SEM analysis further highlighted outer surface softening, wrinkling, and fractures due to water diffusion and plasticization effects. Interestingly, all the other µMESH configurations preserved their architecture over 60 days.

## Conclusion

In this work, the versatility of µMESH is highlighted. Although PLGA is recognized to be a bulk eroding polymer, an increase in SVR of the device did not lead to accelerated degradation, highlighting the complex relationship between these factors. The mechanism was found to strictly depend on the structural dimensions, the incubation medium, and the diffusivity of aqueous media inside the device. Indeed, higher surface-to-volume ratios correlated with slower weight loss and molecular weight reduction in PBS. Among all the tested configurations, the 20×20 µMESH demonstrated the lowest degradation rate, unrevealing a complex interplay between surface-to-volume ratio, polymer amount, and biodegradation.

In conclusion, modifying the microscopic pattern of a polymeric film can regulate its degradation rate and change the dominating mechanism of degradation from a purely bulk one to a combination of bulk and surface erosion.

## Authors’ Contributions

I.G. designed, realized and characterized the µMESH for the in vitro experiments. C.P. realized and characterized the µMESH for the NMR experiment and contributed to the pH analyses and the electron microscopy characterizations. R.S. conducted the in vivo experiments. S.G. and N.T. performed the gel permeation chromatography experiments. D.D.M contributed to μMESH fabrication and characterization. A.L.P. contributed to fluorescence microscopy characterization. P.D. conceived the idea, supervised the whole project, and designed the experiments with I.G.. P.D. and I.G. wrote the manuscript. All the authors have read and contributed to writing the manuscript.

