## Supporting Information for "On the biodegradation of micropatterned polymeric films"

**Supporting Results**

**Visual examination of partially degraded PLGA micronetworks.** PLGAs are known to be hygroscopic polymers [1] and, upon immersion in an aqueous solution, the PLGA micronetwork appeared to soften immediately, suggesting a rapid interaction with the surrounding medium. When PLGA mesh is placed in DI water the concentration of acid groups increases quickly because of its lack in buffer activity, lowering the local pH and boosting the autocatalytic effect. As a result of the plasticizing effect, micronetworks became cloudier and more whitish even after one week, indicating the appearance of little domains of water and a subsequent change in light refractive index. Moreover, specimens appeared rigid and brittle.

When in PBS, the optical and mechanical properties have been maintained for longer as the degradation proceeds more slowly, and the bases partially neutralize the entrapped acidic oligomers. After one week, specimens incubated in PBS did not show any change in color and appeared still ductile. (**Supporting Figure 1A**).

**DI water**

**PBS**

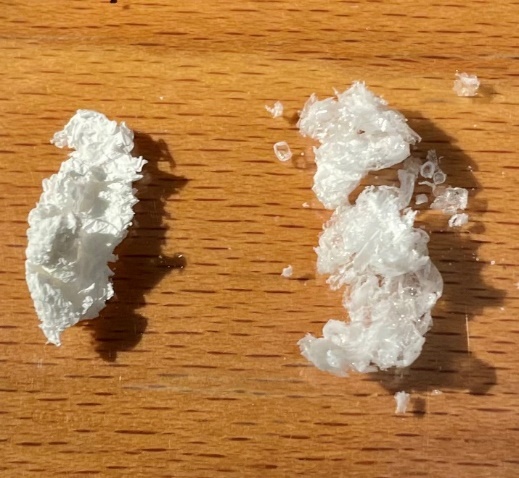

**1 cm**

**Supporting Figure 1**. Images of PLGA micronetworks incubated for 7 days in DI water (***left***) and PBS (***right***). Note that the sample incubated in water appears whitish, whereas the one in PBS appears more transparent and larger in mass.

**µMESH biodegradation in artificial cerebrospinal fluid.** To more accurately recapitulate *in vivo* conditions, µMESH degradation was performed in artificial cerebrospinal fluid (aCSF).

To prepare 500 ml of aCSF, a beaker containing 500 ml of DI water was put on a stir plate at 400 rpm. Then, sodium chloride (105 mM), potassium chloride (2.95 mM), magnesium chloride (4.62 mM), calcium chloride (2.34 mM), sodium carbonate (56.6 mM), disodium hydrogen phosphate dehydrate (0.337 mM), D-glucose (3.33 mM), L-Ascorbic acid (1.14 mM) and bovine serum albumin (0.15 g) were sequentially added and allowed to fully dissolve. Lastly, pH was adjusted to 7.35 ± 0.05 with concentrated hydrochloric acid. All the reagents were purchased from Merck (Darmstadt, Germany). The solution was sterilized using a 0.2 µm pore filter (Corning, supplied by Merck) to avoid bacteria contamination.

Afterwards, 15 mg of PLGA micronetworks presenting 5x5 µm holes were placed into a sterilized sealed jar and incubated in aCSF at 37 ± 0.1°C. At predetermined time points, namely 7, 14, and 21 days, the samples were collected and weighed. The mass loss trend was comparable with the one resulting from degradation in PBS, and in agreement with previous data available in literature [2]. Moreover, no acidification of the medium was recorded. Indeed, PBS and aCSF present similar osmolarity values (~300 mOsm/l [3, 4]) indicating comparable number of salts, which influence pH variation and polymer chain scission. These results are summarized in the plots of **Supporting Figure 2**.

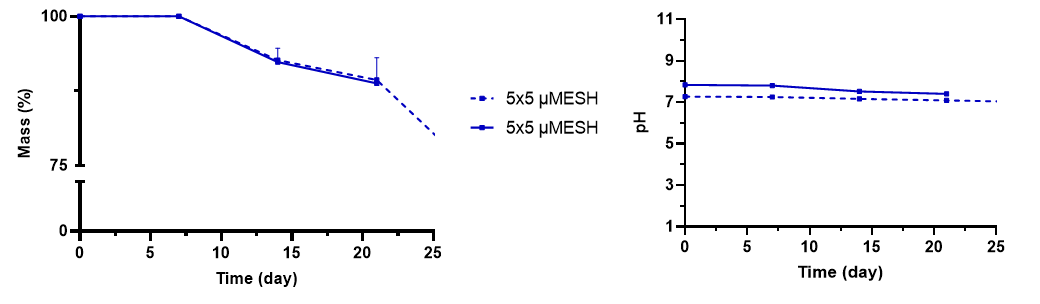

A.

B.

**Supporting Figure 2**. **A** Mass loss vs. time in PBS (dashed line) and aCSF (solid line) for 5×5 µMESH. **B.** Change in pH vs. time in PBS and aCSF for 5×5 µMESH.

**Medium acidification during µMESH** **degradation in a Float-A-Lyzer^®^G2 dialysis device.** Dual-compartment µMESH were placed inside a Float-A-Lyzer^®^G2 dialysis device (cut-off 100 kDa – Fisher Scientific) filled with 2 ml of DI water or 0.1 M PBS pH 7.4. The number of µMESH was adjusted to ensure that all devices contained the same starting amount of polymer (~ 0.5 mg). Then, the dialysis devices were inserted into a 50 ml tube filled with 25 ml of DI water or 0.1 M PBS pH 7.4, according to the medium present inside the device. The tubes were placed into an incubator at 37.0 ± 0.1°C under horizontal rotation at 70 rpm and the pH value of the outer volume was recorded at pre-determined time points. As documented in **Supporting Figure 3**, the pH trends of the incubation media closely resemble those presented in **Figure 2C**, indicating that the PVA microlayer had no significant impact on the acidification of solution.

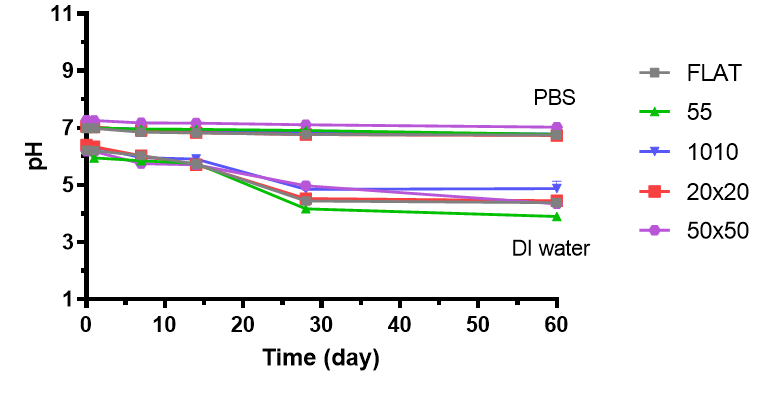

**Supporting Figure 3**. Change in pH vs. time for µMESH in DI water and PBS for all tested configurations and FLAT.

**PULCON method validation from DMSO_2_ samples.** An automated protocol based on the PULCON method was optimized and validated by quantifying gradual concentrations of a certified standard – dimethyl sulfone (DMSO_2_ – 94.13 Da). First, the ^1^H qNMR spectra for dimethyl sulfone (DMSO_2_) standard solutions in deuterated dimethyl sulfoxide ((CD_3_)_2_SO) + 3% TFA were generated with concentrations ranging from 0.5 to 70 mM, using a 400 MHz Avance III spectrometer, at 298 K (**Supporting** **Figure 4A**). Then, the calibration curve (**Supporting** **Figure 4B**), calculated by a least squares regression line, was obtained reporting a strong positive, linear correlation ($y = mc + y_{0}$ with $m$ = 0.9666 and $y_{0}$ = 0.1992 – R^2^ = 0.9991) between the concentrations of DMSO_2_ derived by weighting ($c$) and via the PULCON method ($y$). Whitin the range of 0.5 – 70 mM, the mean relative error and relative standard deviation (RSD) were observed to be smaller than 4% and 5%, respectively (**Supporting** **Figure 4C**). At a lower concentration (0.1 mM), the difference between $c$ and $y$ was found to be 10%, thus, to balance the speed and accuracy of the analysis, a concentration of 0.5 mM was chosen as the Limit of Quantification (LOQ).

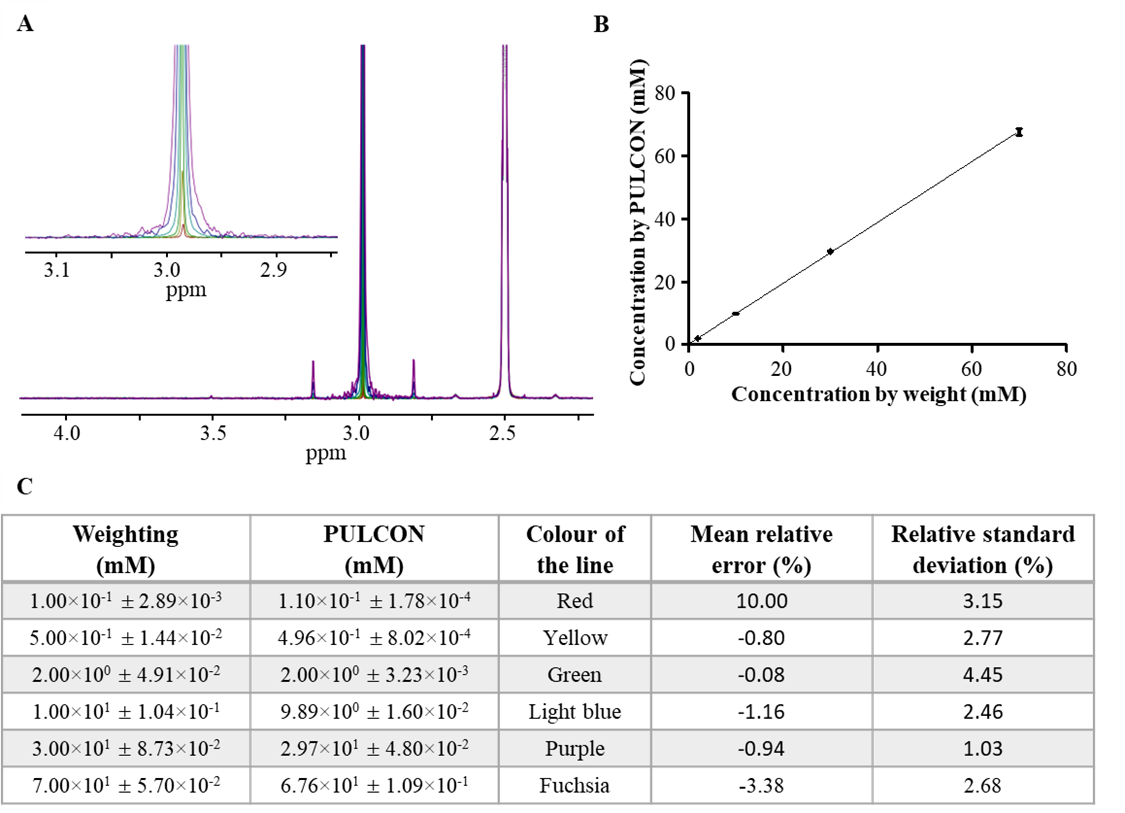

**Supporting Figure 4. Validation of the PULCON method using DMSO_2_ standard samples. A.** ^1^H qNMR superimposed spectra generated on a 400 MHz Avance III spectrometer, at 298 K, for dimethyl sulfone (DMSO_2_) standard solutions in deuterated dimethyl sulfoxide ((CD_3_)_2_SO) + 3% TFA with concentrations ranging from 0.5 to 70 mM. **B.** Calibration curve obtained by plotting the DMSO_2_ concentrations measured by PULCON method (*y*) against those calculated by direct weighting (*c*) within the range of concentration 0.5 –70 mM. A least-squares linear regression method was used to fit the data with the function *y* = 0.9666 *c* + 0.1992 (R^2^ = 0.9991). **C.** A direct comparison between the concentrations of DMSO_2_ obtained by weigh (first column) and measured by PULCON (second column). A color legend listed in the third column reports the color of the line in the superimposed spectra per DMSO_2_ concentrations. The fourth and fifth columns list the mean relative error and relative standard deviation, respectively. Data are reported as mean ± absolute uncertainty for n = 3 independent repetitions. The larger error occurs at the lowest concentration of DMSO_2_ (0.1 mM).

**PULCON method for PLGA mass quantification.** The PULCON method was validated for poly(lactide-co-glycolyde) (PLGA) quantification following a partially modified protocol previously described by the authors [5]. Samples at three concentrations of PLGA, namely 2.84, 28.4 and 141 µM (repetitive unit concentrations for a 38 – 54 kDa PLGA: 1, 10 and 50 mM, respectively), were prepared in triplicate in ((CD_3_)_2_SO) + 3% TFA and analyzed by using a Bruker Avance III 400 MHz spectrometer. Both the signals of methine (-CH) group at 5.05 – 5.30 ppm and methyl (-CH_3_) group at 1.35 – 1.55 ppm of PLGA were considered for the quantification (**Supporting** **Figure 5A**). A strong correlation was obtained between the concentrations obtained by weight (first column) and concentrations calculated by PULCON for both the methine and methyl signal (second and fourth column, respectively), showing a mean relative error lower than 10% (**Supporting** **Figure 5B**).

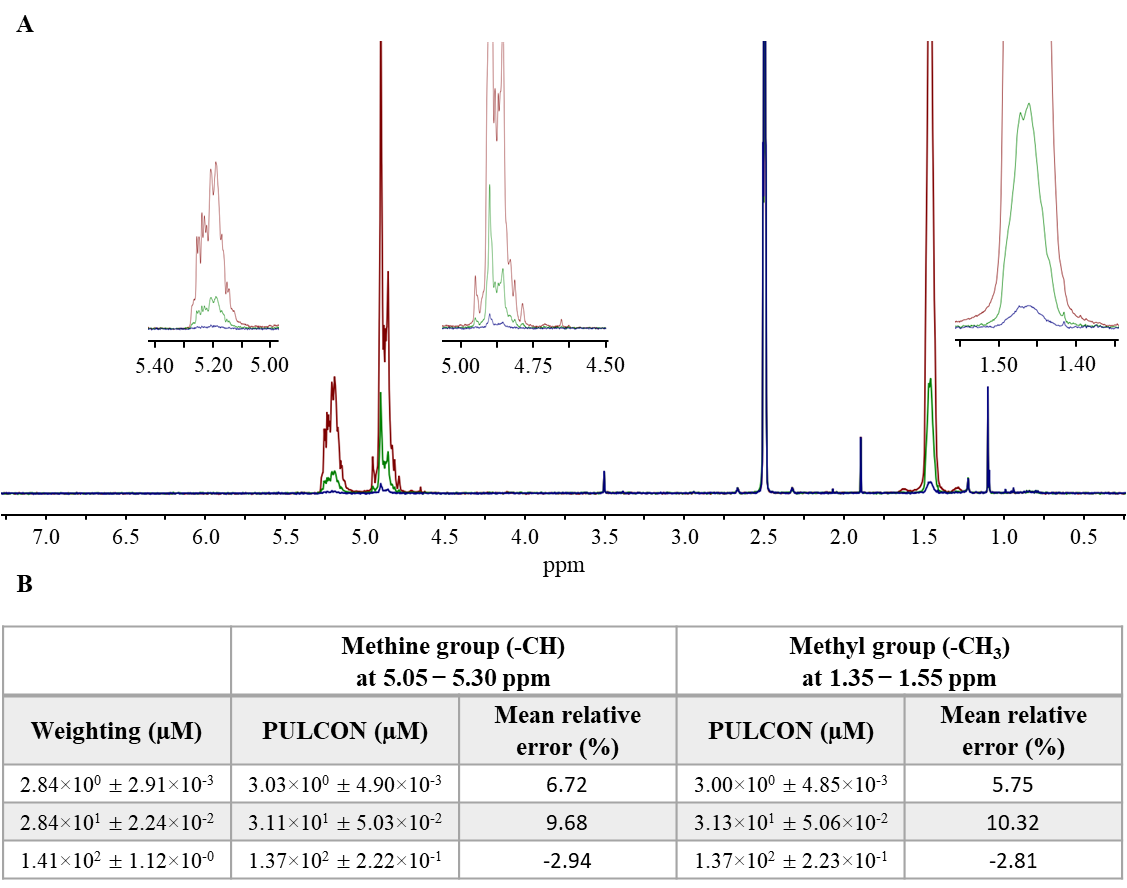

**Supporting Figure 5. A.** ^1^H qNMR superimposed spectra generated on a 400 MHz Avance III spectrometer, at 298 K, for PLGA solutions in deuterated dimethyl sulfoxide ((CD_3_)_2_SO). **B.** A direct comparison between concentrations of PLGA obtained by weight (first column) and measured by PULCON considering the methine group (second column) and considering the methyl group (fourth column). The third and fifth columns list the mean relative errors of data measured by PULCON on methine and methyl groups, respectively. Data are reported as mean ± absolute uncertainty for n = 3 independent repetitions).
